# Strategy for selective acquisition of transgenic marmosets using the piggyBac transposon system

**DOI:** 10.1101/2022.08.02.502441

**Authors:** Tsukasa Takahashi, Takuji Maeda, Motohito Goto, Masafumi Yamamoto, Wakako Kumita, Ryoji Ito, Tomoo Eto, Takashi Inoue, Hideyuki Okano, Erika Sasaki

## Abstract

Transgenic (Tg) nonhuman primates (NHPs) are excellent potential models due to their physiological similarities with humans. In NHPs, the lentiviral vector is the only available transgenesis method, but transgene sizes are limited. The piggyBac (PB) transposon system is a DNA element sandwiched between two terminal inverted repeats (TRs) that allows transposases to introduce larger genes than lentiviral vectors. However, embryos with integrated transgenes are difficult to distinguish using this system. To address this issue, we placed the Enhanced Green Fluorescent Protein (EGFP) gene inside the TRs and the humanized kusabira orange 1 (hKO1) fluorescent protein gene with the proteolysis-promoting sequence d2PEST outside the TRs to create an indicator for gene integration using fluorescent protein expression. In HEK293T cells, hKO1 and EGFP co-expression after transfection gradually changed to decay of hKO1 expression and enhancement of EGFP expression by transposase within five days. In mouse and marmoset embryos, hKO1 expression was attenuated, while only EGFP expression was observed at the blastocyst stage. All offspring obtained through embryo transfer of EGFP-expressing embryos were Tg; thus, we established a new PB system capable of determining Tg embryos and successfully produced Tg marmosets, for the first time, in NHPs.

## Introduction

The marmoset (*Callithrix jacchus*) is a small new world primate used in biomedical research owing to its physiological similarities to humans, small body size, and fecundity. The marmoset’s prolificacy is the most important biological feature for producing genetically modified marmosets. Marmosets reach sexual maturity in 12 to 18 months, earlier than other nonhuman primates (NHPs). They ovulate two or three oocytes in each ovarian cycle and produce offspring at approximately 155 days; a single female can produce 4–6 offspring per year. Thus, the marmoset is among the most fecund NHPs and is a suitable experimental animal for genetic modification^1-4^.

The first Tg NHPs generated were marmosets produced using the lentiviral vector method and successfully passed on transgenes to the next generation^5^. The lentiviral vector technology is the most efficient transgenesis method and has been used in rodents and various farm animals^6,7^. The use of lentiviral vectors has made it possible to produce Tg animals, even in NHPs, where it is difficult to obtain a large number of oocytes ^2,8-10^. A transgene undergoes protein expression when inserted into the host genome using a lentiviral vector, allowing for the selection of tg embryos based on whether the reporter gene is expressed for embryo transfer to surrogate mothers^11^. This embryo selection results in all Tg animals offspring^5,9^, and is also a suitable method for NHPs to avoid euthanasia of failed Tg animals owing to ethical reasons. However, lentiviral vectors have disadvantages such as reduced pregnancy rates due to embryotoxicity^12^ and a transgene size limit of up to 8 kb^13^. Particularly, the limited transgene-carrying capacity prevents the generation of various disease models.

DNA transposon systems have been applied for advancing the production of Tg animals in many species^12,14-19^. DNA transposon is a genetic element that mobilizes DNA sequences using transposase. The transposase binds to terminal inverted repeats (TRs), which are recognition sites at both ends of the DNA sequence, excises the DNA segment flanked by TR from the genome, and pastes the segment to a new location. One of the transposons, *piggyBac* (PB), has been isolated from the cabbage looper moth (*Trichoplusia ni)*, and proven to be the most efficient transposon in mammalian cells^20-22^. Hyperactive PBase (hyPBase), which shows a higher transplantation rate than conventional Wild-type (WT) mammalian codon-optimized PBase, has also been developed and applied in Tg animal production^23^. They have high transduction efficiency and can successfully perform long gene transfer^12,14^. Furthermore, PB can introduce genes into the zygote by microinjection into the pronucleus as well as intracytoplasmic sperm injection transgenesis (ICSI-Tr) and intracytoplasmic injection transgenesis^14^. Therefore, genetically modified animals have been produced by PB even in animal species in which microinjection of the pronucleus is difficult owing to the color or structure of fertilized oocytes^24^. However, data on the production of genetically modified animals by PB in NHPs are lacking. Additionally, since PB injects plasmid DNA, distinguishing whether the expression of the reporter gene originates from the episome or gene integrated into the genome is difficult, which hinders the application of PB in generating Tg NHPs, including marmosets.

We conducted this study to develop a new PB vector system that can determine the integration of a transgene into the host genome by expressing two reporter genes. PB has a unique characteristic: only the DNA sequence flanked by TRs is inserted into the host genome. Using this characteristic, we placed two different fluorescent proteins inside and outside the TR of PB vectors change fluorescent protein expression patterns upon transgene integration into the host genome. Furthermore, using the PB system, we generated the first Tg NHPs and showed that the embryo selection via fluorescent protein expression is highly efficient for Tg animal production.

## Results

### *PB* vectors with two fluorescence proteins for determining gene integration

This study designed a new PB vector with two differently colored fluorescence proteins for determining transgene integration. A schematic diagram of the PB vector is shown in Fig. 1a. The vector contains a green fluorescent protein (EGFP) gene under the CMV promoter between the TR sequence and the humanized-Kusabira-Orange 1 (hKO1) gene with the proteolytic sequence d2PEST under the CMV promoter outside the TR sequence. A transposase, hyPBase, recognizes the TR sequence, cuts out the CMV-EGFP gene sequence, and pastes it into the host genome; CMV-hKO1d2PEST does not integrate into the host genome. As a result, only EGFP expression continues, and hKO1 expression gradually decreases. Changes in the two colors due to changes in the expression of the two fluorescent proteins can determine the transgene integration into the chromosome. To prove this concept, the plasmid PB-CMV-hKOd2PEST-SVpA-CMVp-eGFP-pA (PB) and/or the hyperactive PBase expression vector pCMV-hyperactive PBase (hyPBase) were transfected into HEK293T cells. Changes in EGFP and hKO1 expression were observed for 15 days to determine the timing of transgene integration (Fig. 1b). On Day 1, 39.6% and 32.0% of the cells co-expressed EGFP and hKO1 in the PB-alone and PB+hyPBase groups, respectively (Fig. 1b, i, iv-vi, xiii, xvi-xviii). On Day 5, fluorescence-activated cell sorting (FACS) analysis showed that 18.9% of the cells were double-positive in the PB-alone group, whereas 0.67% expressed EGFP alone (Fig. 1b, ii, vii-ix). Additionally, in the PB+hyPBase group, 17.1% cells co-expressed EGFP and hKO1, whereas 10.7% expressed only EGFP (Fig. 1b, xiv, xix-xxi). On Day 15, only 0.49% of EGFP-positive cells were present in the PB-alone group, whereas 15.6% of EGFP-positive cells were detected in the PB+hyPBase group (Fig. 1b, iii, x-xii, xv, xii-xxiv). The PB-alone group showed changes in the transient expression of fluorescent proteins. In contrast, the PB+hyPBase group showed continuous changes in fluorescent protein expression owing to active integration. The increasing number of EGFP expressing cells indicated that hyPBase incorporated the CMV-EGFP sequence between TRs into the host chromosome in the PB+hyPBase group.

**Fig. 1.**
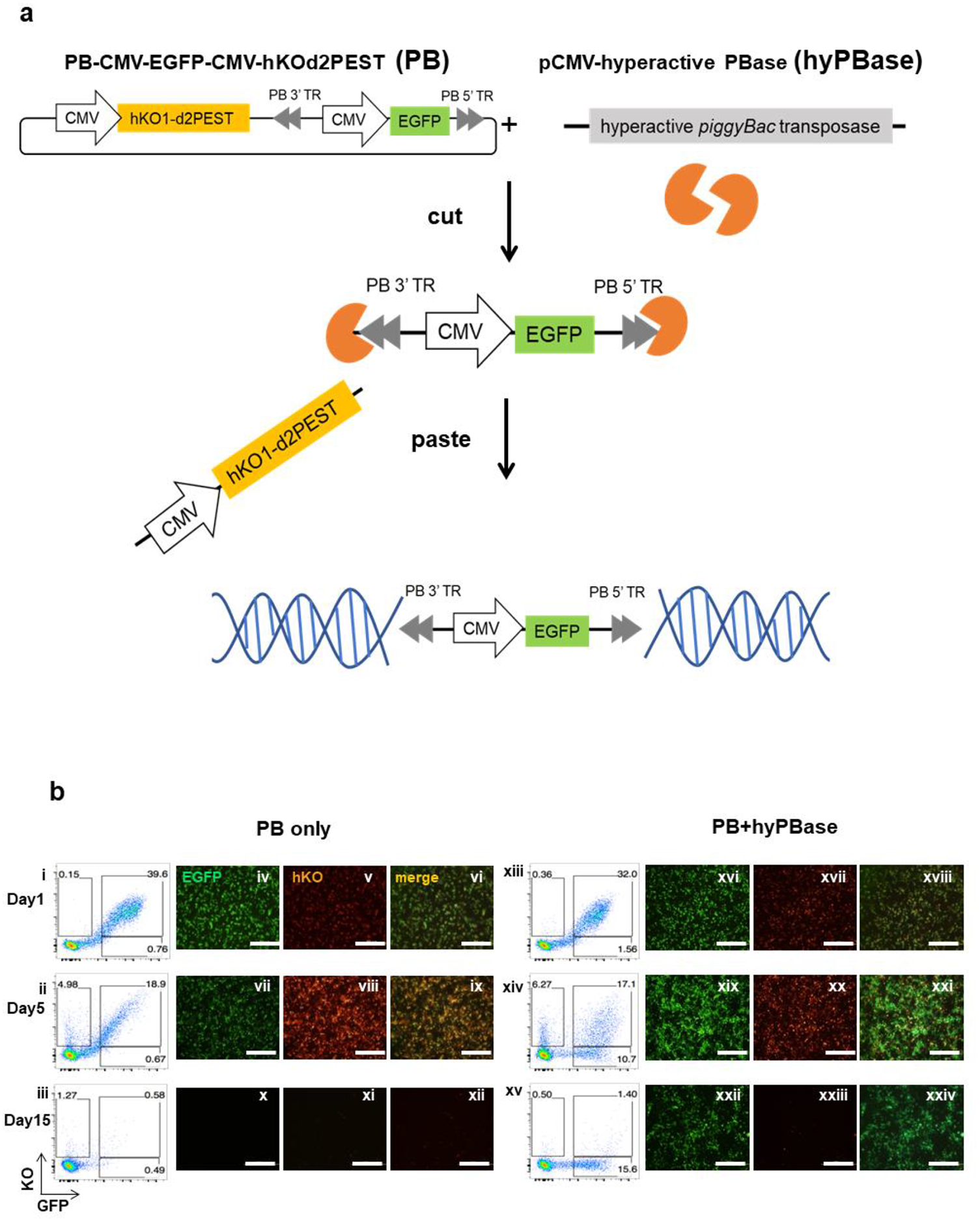
(a) Schematic diagram of piggyBac (PB) vector with selection markers. A hyperactive piggyBac transposase (hyPBase) cuts only CMV-EGFP in the TR and pastes it to the chromosome while CMV-hKOd2PEST outside the TR is cut off and attenuated. (b) Fluorescence expression and FACS analysis of HEK293T cells transfected with PB vector alone or PB vector and hyPBase. (i)-(iii) and (xiii)-(xv) are images of FACS analysis. (iv)-(xii), (xvi)-(xxiv) are images of EGFP and hKO1 merged on each day. Scale bar = 200 μm

### Tg embryo selection following PB vector injection in mice embryos

To determine whether it was possible to select Tg embryos in mice, we microinjected 2 ng/μL of PB-alone or 2 ng/μL of PB and 1ng/μL of hyPBase in the pronucleus of zygotes. We observed EGFP and hKO1 expression during development (Fig. S1). On Day 1 post-injection, in both PB-alone and PB+hyPBase groups, EGFP and hKO1 expression were observed in some embryos (Fig. S1a). On Days 2-4, in the PB+hyPBase group, the number of embryos expressing only EGFP gradually increased, whereas the PB-alone group showed co-expression of EGFP and hKO1. Further, a single embryo was analyzed for 4 days to analyze changes in fluorescent protein expression patterns. Unlike in HEK293T cells, in mouse embryos, hKO1 expression could not be detected from the first day till the end (Fig. S1b).

To validate this selection criterion, we microinjected 0.67 ng/μL of PB and 0.33 ng/μL of hyPBase into the pronucleus, selected blastocysts expressing only EGFP on Day 4 for embryo transfer and evaluated the birth rates of Tg progeny (Table S1). When the transplanted embryos were selected without considering fluorescent protein color, 50% of the offspring were Tg mice; while when the transferred embryos were selected by evaluating EGFP expression, 100% of the neonates were Tg mice (Table S1).

Cytoplasmic injection of mice embryos was performed using 2 ng/μL of PB and 1 ng/μL of hyPBase (Table S1). When embryos were transplanted without evaluating fluorescent protein color, 81.8% of offspring were Tg animals, and when blastocysts expressing only EGFP were selected for embryo transfer, 100% of neonates were Tg mice (Fig. S2a). These results demonstrated that the selection of transferring embryos using fluorescent proteins encoding in TR sequences was accurate. Further, with this selection, the PB vector could produce only Tg mice. The F0 mice were bred with WT, and progeny tests showed both nuclear and cytoplasmic injection of PB+hyPBase, resulting in germline transmission (Fig. S2b, Table S2.). FISH analysis of three F1 mice (one with nuclear and two with cytoplasmic injections) showed gene transfer at 1–2 chromosomal sites (Fig. S2b).

One Tg mouse with hKO1 outside TR was also obtained in both pronuclear and cytoplasmic injection groups, and hKO1 was also inherited in F1 progeny.

### Selective acquisition of Tg marmosets using PB vectors

Microinjection was performed in marmoset embryos to apply this PB vector-based genetic manipulation. In marmosets, a cytoplasmic injection would be more effective than a pronuclear injection since the marmoset pronuclei of fertilized oocytes are small. Therefore, 1 ng/μL of hyPBase and 2 ng/μL of PB or 2 ng/μL of PB alone were injected into the cytoplasm of marmoset zygotes and fluorescent protein expression was observed (Table 1). In the PB+hyPBase microinjected groups, 44 of the 56 embryos grew to the 2-cell stage. On day 4 after injection with PB+hyPBase, 72.7% (32/44) of the embryos expressed fluorescent proteins hKO1 and/or EGFP, indicating successful injection. Of these, 34.6% (11/32) embryos expressed both hKO1 and EGFP, and 65.6% (21/32) embryos expressed only EGFP (Table 1 and Fig. 2 a, b). Furthermore, by Day 7, 59.4% of the embryos expressing fluorescence continued to develop into morula and blastocysts. Of these, 84.2% (16/19) expressed only EGFP, and hKO1 expression was completely undetectable via fluorescence microscopy (Table 1 and Fig. 2a). In contrast, 15.8% (3/19) of the embryos strongly expressed hKO1 without any decay (Fig. 2b).

**Table 1.**
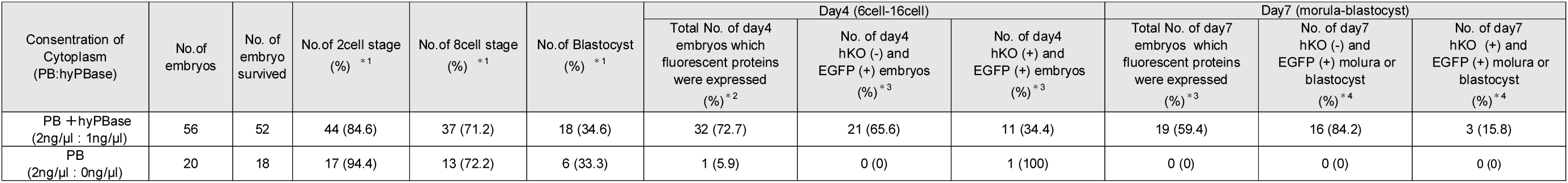
Embryo development and fluorescence protein expression rates after PB cytoplasmic injection in marmosets. *1: number in parentheses was calculated from total survived embryos. *2: number in parentheses was calculated from 2-cell. *3: number in parentheses were calculated from the total no. of Day 4 embryos witch fluorescent protein expression. *4: number in parentheses was calculated from total no. of Day 7 embryos with fluorescent protein expression.

**Fig. 2.**
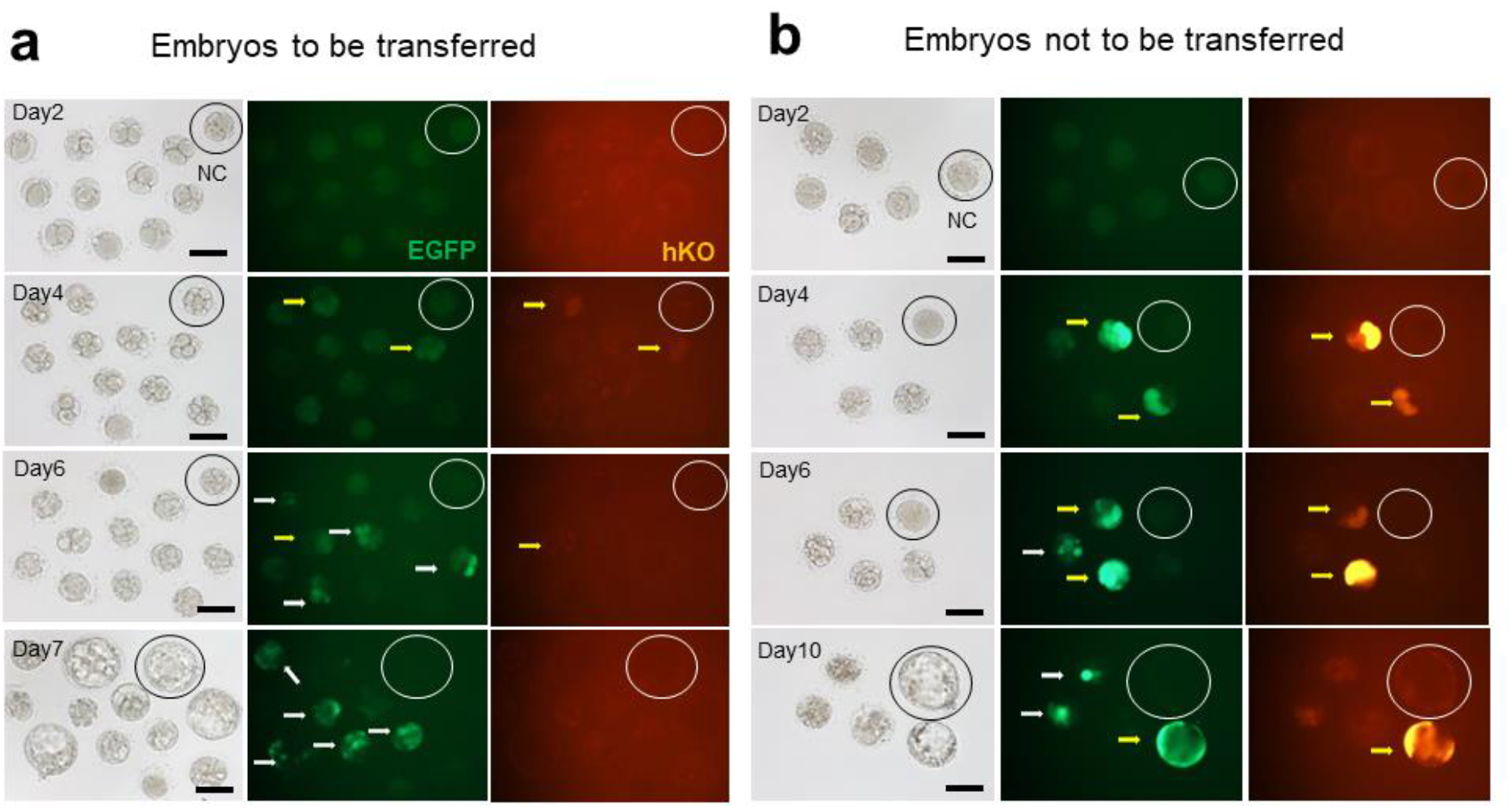
Fluorescence protein expression changes during embryogenesis after microinjection of PB and hyPBase into the cytoplasm of marmoset fertilized oocytes. Yellow arrows indicate embryos with co-expression of hKO1 and EGFP. White arrows indicate embryos expressing only EGFP. (a) Embryos to be transferred: As embryogenesis progresses, only EGFP was expressed (white arrow). (b) Embryos not to be transferred: hKO1 does not decay during development, and hKO1 and EGFP remain co-expressed (yellow arrows) until the blastocyst stage. Circles indicate negative control embryo. Scale bar = 100 μm.

In the PB-alone group, hKO1 and EGFP co-expression was observed in 5.9% (1/17) of Day 4 embryos hKO1 expression did not decrease the expression level. On Day 7, all six blastocyst embryos did not express hKO1 or EGFP (Table 1).

While mouse embryos develop blastocysts within 4 days, the *in vitro* developed marmoset embryos take approximately 7–10 days to reach the blastocyst stage. The expression of fluorescent proteins in marmoset embryos was observed after day 4 of the PB+hyPBase microinjection (Fig. 2a, b). Based on these results, marmoset embryos that strongly expressed EGFP after Day 4 were considered Tg embryos and were used for embryo transfer when hKO1 fluorescence intensity decreased.

Nineteen EGFP-expressing embryos developed to the morula or blastocyst stage after 6– 7 days of culture and were transferred to 12 recipient female animals for producing tg marmosets. Of these, 4 (33.3%) were pregnant, and all miscarried (Table 2). On the other hand, following embryo transfer of 30 EGFP-expressing embryos at the 6-to 16-cell stage to 19 recipient animals, three (15.8%) animals became pregnant, and one gave birth to twins (Table 2, Fig. 3a).

**Table 2.** In vivo development of EGFP-expressing embryos after PB and hyPBase cytoplasmic injection in marmosets.

| day after injection | stage of embryo | No. of embryo transferd | No. of recipient | No. of pregnant (%recipient) | No. of miscarriages (%pregnant) | No. of offspring (% embryo) | No. of EGFP ( + )<br>Tg offspring (% offspring) | No. of hKO ( + )<br>Tg offspring (% offspring) |
| --- | --- | --- | --- | --- | --- | --- | --- | --- |
| Day6-7 | Morula-Blastocyst | 19 | 12 | 4<br>(33.3) | 4<br>(100) | 0<br>(0) | 0<br>(0) | 0<br>(0) |
| Day4-5 | 6cell-16cell | 30 | 19 | 3<br>(15.8) | 2<br>(66.7) | 2<br>(6.7) | 2<br>(100) | 0<br>(0) |

**Fig. 3.**
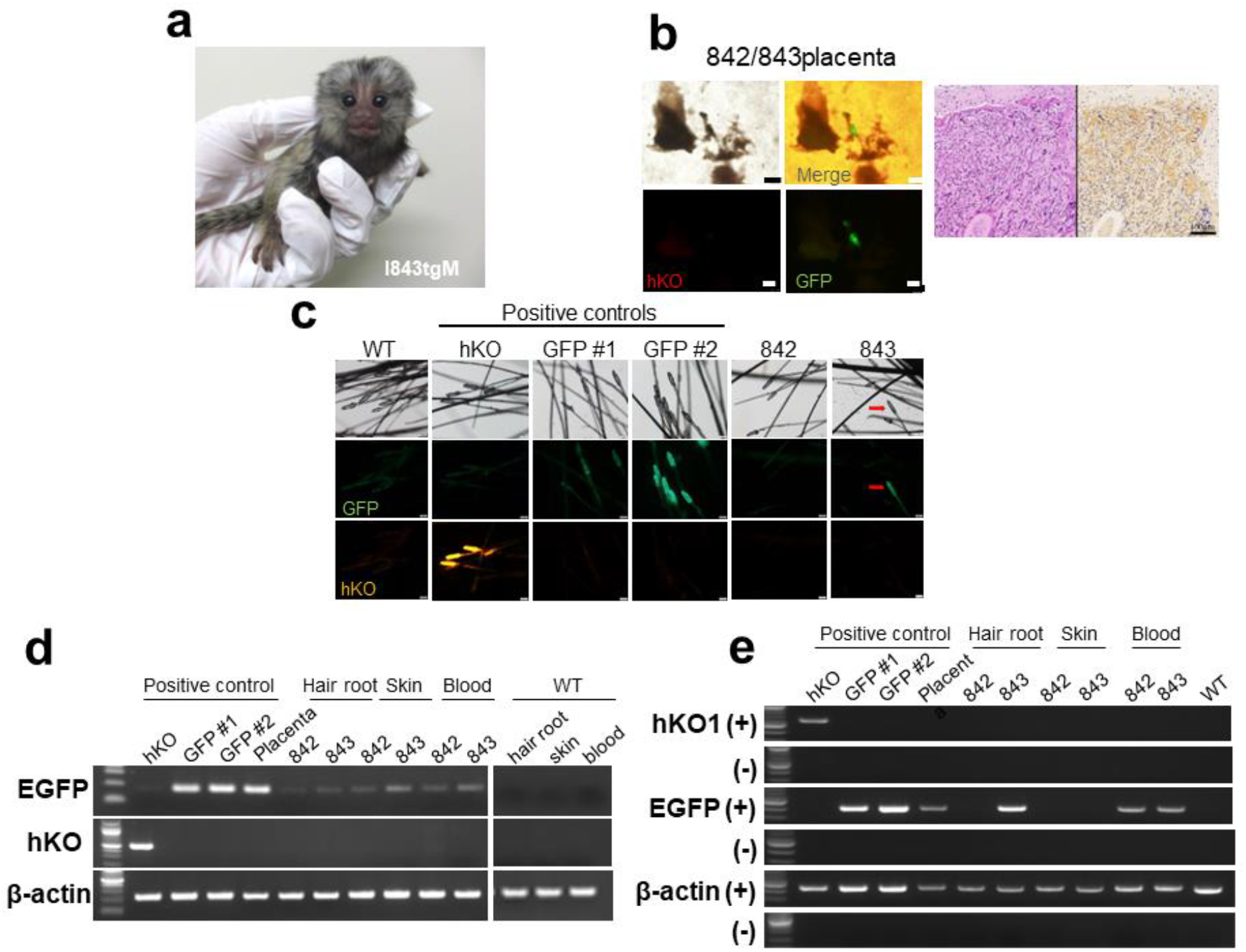
Genotyping of offspring. (a) a photograph of the offspring #843. (b) Placenta of offspring #842 and #843. Fluorescent protein images and immunostaining of EGFP in tissue sections. (c) Fluorescence images of hair root. EGFP PC#1 is a one-copy transgene with weak expression. PC#2 is a one-copy transgene with strong expression. Red arrow: hair root of expressing EGFP. (d) Genomic PCR of the hair root, skin, and blood of offspring. (e) RT-PCR of the hair root, skin, and blood of offspring. Scale bar = 100 μm

### Genotyping of newborn marmosets

Since marmoset twins share a placenta by fusion of their own placentas^25-28^, only one placenta was obtained from the twin neonates. Fluorescence microscopy images showed EGFP expression but not hKO1 in parts of the neonates’ placenta; immunostaining of the placenta also detected EGFP (Fig. 3b).

Next, we compared the fluorescence images of hair roots with the positive controls (PCs) of hKO1 Tg and EGFP Tg marmosets. The PC hKO1 Tg marmoset expressed a high level of hKO1 and carried more than 10 copies of the hKO1 gene (data not shown). The PC of Tg marmoset, EGFP PC #1, expressed low levels of EGFP and carried a single copy of the EGFP gene, and EGFP PC #2 expressed high levels of EGFP and carried a single copy of the EGFP gene. As a result, hair root fluorescence imaging detected strong EGFP expression in some hair roots in #843; however, no EGFP expression was observed in the hair roots of #842 (Fig. 3c).

PCR and RT-PCR were performed using DNA and mRNA collected from neonates’ hair roots, skin, and blood samples (Figs. 3d, 3e, S3, and S4). Although PCR using genomic DNA showed that both animals carried the EGFP gene, the PCR band intensities were much lower than those of the single-copy positive control. RT-PCR results revealed that EGFP mRNA transcriptions were observed in the placenta, blood sample of animal #842, and hair roots and blood of animal #843 but not in the skin samples of both neonates. On the other hand, hKO1 mRNA transcription was not observed in all tested samples.

After sexual maturation of #842 and #843, *in vitro* fertilization was performed using #843 sperm. The results showed that 1 out of 31 fertilized oocytes (3.2%) showed EGFP expression at the 8-cell stage, and 1 out of 82 embryos (1.2%) collected from naturally mated oocytes of females paired with #842 showed EGFP expression at the blastocyst stage (Table 3 and Fig. 4).

**Table 3.** Evaluation of germline transmission by generating EGFP-positive fertilized oocytes using sperms from male founder animals.

| Animal number | Method of embryo collection | No. of obtained Fertilized oocytes | No. of EGFP (+) embryo (%) |
| --- | --- | --- | --- |
| 843 | IVF | 31 | 1 (3.2) |
| 842 | Natural mating | 82 | 1 (1.2) |

**Fig. 4.**
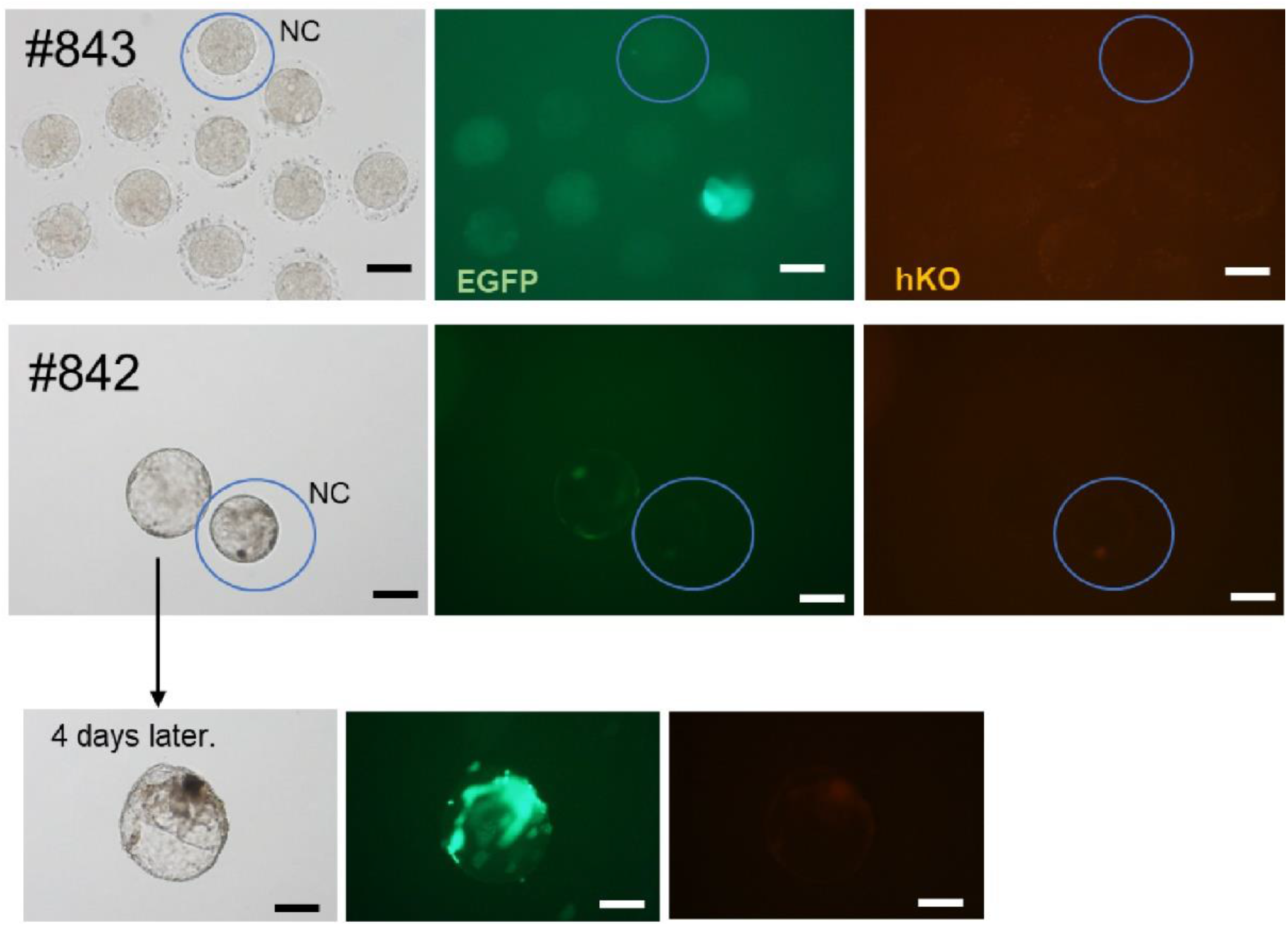
EGFP-positive embryos fertilized with sperms from male animals. Top panel: 8-cell stage of IVF embryo #843. Middle panel: Blastocyst stage of naturally mated oocyte of #842. Bottom panel: EGFP became stronger by culturing the naturally mated oocyte of #842 in the middle panel for another 4 days. Scale bar = 100 μm.

## Discussion

In this study, we developed a new PB system that could visualize transgene integration into the host genome, and we successfully produced the first Tg NHP using the PB system. In the PB system, transgene integrations were visualized by changes in fluorescent protein expression patterns. Specifically, during cell/embryo culture, decreasing hKO1 expression encoded outside TR and continuing EGFP expression encoded inside TR indicated transgene incorporation into the host genome. Therefore, for the generation of Tg NHPs, the PB system allows for the selection of Tg embryos before embryo transfer and obtain Tg marmosets with a 100% success rate. This PB system can reduce the number of produced animals and avoid euthanasia of animals that fail to integrate transgenes into the host genome. Particularly, primates have a long lifespan and high maintenance costs, and from the perspective of animal welfare, it is essential to ensure not to produce failed genetic manipulation animals. Our PB system is a suitable method that complies with the “reduction” and “refinement” of the 3Rs of guiding principles for more ethical animal use in laboratory testing.

In transfection of HEK293T cells, hKO1 and EGFP were co-expressed on Day 1 with or without PBase. On day 5, the FACS analysis showed that the PB and PBase group had more cells expressing only EGFP but, not in the PB-alone group (Fig. 1b). Fluorescent microscopies also showed a clear difference, as illustrated in Fig. 1b-(ix) and (xxi), suggesting that the switch in fluorescent protein expression within 5 days can be an indicator for gene integration. By day 15, cells in which the gene was inserted entirely switched to EGFP single expression.

Cells can be cultured for a long time, and changes can be seen until the last phenomenon. However, mouse embryos reach the blastocyst stage 4 days of culture, while marmosets reach the blastocyst stage at 7 to 10 days of culture, making it necessary to make decisions in a short period. Additionally, the long-term culture of embryos decreases pregnancy and birth rates^29^, and thus, it is essential to determine gene transfer in as less time as possible. Comparing the PB+hyPBase microinjected into mouse embryos and the PB-alone group, the merged image of the blastocysts on day 4 (Fig. S1a) was similar to the HEK293T (Fig.1b-(ix), (xxi)). Therefore, the criteria for transferring embryos to surrogate mothers were weak or no expression of hKO1, and robust EGFP expression were observed, on Day 4. The embryo transfer results of both mouse and marmoset showed that all offspring were transgenic neonates, indicating that our criteria can select the transgene integrated embryos.

On the other hand, the embryos with continuing strong expression of hKO1 and co-expression of EGFP after day 4 were judged as transgene-non-integrated embryos, however, rarely found (Table S1).

As a result of transferring mouse embryos without sorting EGFP expression, 50% of Tg animals were obtained in pronuclear injection and 81.8% in cytoplasmic injection, suggesting that with microinjection of PB+hyPBase, a high rate of successful gene integration occurred. The attenuation of hKO1 outside TRs was clear in cells but unclear in embryos. In mouse and marmoset embryos, hKO1 expression was weak or absent, and only EGFP expression was elevated in some embryos (Figs. 2 and S1). In this study, the hKO1 connected to d2PEST, a proteolysis-promoting sequence^30^. For HEK293T cells evaluation, d2PEST was selected since it shows the most precise color switch to EGFP compared to that without the PEST sequence or d4PEST with its long half-life ^31^.

The expression of fluorescent proteins in embryos was weak because the copy numbers of genes introduced into the embryos via microinjection might be less than those of genes introduced into cells via lipofection. The expression of fluorescent proteins in the embryo weakened. This can be solved by adjusting the concentration of embryo injection or switching to a reporter with high detection sensitivity. The expression of EGFP in TRs increased from Day 1 in mouse embryos, whereas it increased from Day 4 in marmoset embryos. The timing of gene expression in the embryo is also related to the Zygotic Gene Activation (ZGA), which is different from that of other species. ZGA suggests that developmental progression during the first few days after fertilization is controlled by maternal mRNA, and then switches to regulation by embryo-derived transcripts. In mice, the embryo-derived transcriptional activity is greatly increased at the 2-cell stage^32-34^, while in humans, it occurs in 4 to 8-cell stage embryos, and in cattle, in 8 to 16-cell stage embryos^35 36^. Marmoset ZGA occurs at a similar stage as that of human embryos^37^. Therefore, the timing of genome activation varies among animal species^35^. Therefore, the evaluation of changes in fluorescence expression in embryos should be designed considering the genome activation time.

In this study, we obtained Tg mice carrying the hKO1 gene inserted outside TR sequences. PB vectors were injected as a circular plasmid DNA, and genomic PCR detected outer frames of PB vectors, including hKO1, suggesting that the vector backbone flanked by outer TRs was recognized by hyPBase and introduced into the host chromosome. This can be overcome by linearizing the plasmid and not sandwiching a negative selection marker, hKO1 in this case, between TRs. However, linearization of transposons inhibits embryonic development^19^. When we compared the injection of circular and linearized plasmid DNA, we found that the rate of development of blastocyst was three times higher in circular DNA (data not shown). Thus, although the toxicity of linearized plasmid DNA was not clarified, it is necessary to optimize the injection conditions when linearized PB injection is required to avoid interference with embryonic development.

Lentiviral vectors are highly embryotoxic and have a very low pregnancy rate^12^. While the pregnancy rate of marmoset embryos injected with PB was high (15.8–33.3%) and comparable to untreated WT embryos^29^, the miscarriage rate was very high (66.7–100%; Table 2), with a similar offspring birth rate to that of lentivirus vectors transgenesis^5^. Although PB integrated encoded genes between the TR and target site of PB, known as the TTAA sequence^20^, the transgene was randomly integrated and the copy number of the gene could not be controlled. Therefore, embryonic lethality may occur when transgene integration disrupts a major gene.

The two Tg marmosets showed a high degree of mosaicism. Mosaic transgenesis via transposon has been previously reported ^38,39^. To prevent embryo mosaicism, hyPBase with mRNA was injected for swift expression and rapid decay. However, mosaicism could not be prevented, uninformed expression of EGFP was observed in embryos under fluorescence microscopy, and mosaicism was observed for expression in tissues. Particularly, since the PB vector and hyPBase were injected into the cytoplasm of marmoset embryos, it took a long time for gene integration than for pronuclear injection, which might have contributed to the increased mosaicism rate. To prevent mosaicism, it is crucial to express hyPBase at an early stage. It is essential to develop new technologies such as ICSI-Tr^14^ to inject nucleic acids into the oocytes or new methods for swiftly expressing PBase protein to solve the mosaic rate issue. Although relatively strong expression was observed in some embryos and hair roots of #843, no expression was detected in the fibroblasts and blood cells via FACS analysis. It has been reported that the promoter of CMV is generally active throughout the body; however, it is tissue-specific in marmosets^5^. This characteristic of the promoter led to variations in the expression of each tissue.

In this study, we developed an efficient PB system that could select Tg embryos and obtained the first transposon mediated Tg offspring in NHPs. The PB system was highly efficient in gene transfer and enabled larger gene transfer than lentiviral vectors or knock-in technology. In the future, the PB system will expand the possibility of creating disease models in NHPs.

## Methods

### Animals

All animal experiments were approved by the Animal Committee of the Central Institute for Experimental Animals (approval numbers: 11028A, 14029A, 15020A, 16019A, 17029A, 18031A,19033A, 20049A) and performed according to CIEA guidelines, which are by the guidelines for “The proper conduct of animal experiments” determined by the Science Council of Japan. C57BL/6JJcl and B6D2F1/Jcl mice were used to collect zygotes at the pronuclear stage (female: 8–16 weeks old, male: 12–20 weeks old). Jcl:MCH (ICR) female mice aged 10–16 weeks were used as recipients for embryo transfer. All mice were purchased from CLEA Japan Inc. (Tokyo, Japan) and reared under the following conditions: room temperature, 22°C ± 0.5°C, humidity, 55% ± 5%; lights on 08:00–20:00. Food (CA-1; CLEA Japan Inc.) and water were provided *ad libitum*.

The marmosets were housed in pairs in stainless steel living cages (W820 x D610 x H1578 mm) with wire mesh floors maintained at 25–27°C with 45–55% humidity and illumination for 12 h per day. Wood perches for locomotion and gouging and a platform, a nest box, or a hamockfor bed were placed in each cage for environmental enrichment. Marmosets were kept healthy and well-nourished with a balanced diet (CMS-1M; CLEA Japan Inc.), including mixed L (+)-ascorbic acid (Nacalai Tesque, Tokyo, Japan); vitamins A, D3, and E (Duphasol AE3D; Kyoritsu Seiyaku Co., Ltd., Tokyo, Japan); and honey (Nihonhatimitsu Co., Ltd., Gifu, Japan). Additionally, chicken liver boiled in water (DBF Pet Co., Ltd., Niigata, Japan) was given as a supporting meal weekly. The animals were supplied with tap water *ad libitum* from feed valves. All animals were at least 2 years old and weighed 300 g or more. They were all purchased from a marmoset breeding company for experimental animals (CLEA Japan Inc.). Marmoset oocytes and natural mating embryos were collected from 33 sexually mature females. Sperms were collected from eight sexually mature males, and 38 recipients were used.

### PB vector construction and hyPBase mRNA production

The PB vector (PB-CMV-hKOd2PEST-SVpA-CMVp-eGFP-pA) was constructed as described below. The following procedure was used for hKO1 fluorescence to be placed in the TR outer side of the PB vector. The hKO1d2PEST fragment was cut from pENTR-hKOd2PEST (kindly provided by Dr. Kanki, Keio University) using BamHI/EcoRI and blunting. pmKate2-N (evrogen, Moscow, Russia) was cut using NheI/HpaI to remove mKate. The hKO1d2PEST fragment was ligated next to the pCMV of pmKate2-N to produce pCMV-hKO1d2PEST. This pCMV-hKO1d2PEST fragment was cut out with AflII/AflIII and blunting. PB-R4R2-DEST, the backbone of the PB vector was cut and blunted with KpnI, and the pCMV-hKO1d2PEST fragment was ligated to the blunted part to construct PB-CMV-hKOd2PEST-SVpA -R4R2-DEST. pCMV-EGFP to be placed in the inner side of the TR of PB vector was constructed as follows: pENTR5’/CMVp was purchased (Thermo Fisher Scientific Inc. Waltham, MA), and EGFP cDNA was cloned into pENTR1A (Thermo Fisher Scientific Inc.) vector. PB vectors were constructed using LR plus reaction (Thermo Fisher Scientific Inc.) of pCMV entry vector, EGFP entry vector, and PB-CMV-hKOd2PEST-SVpA-R4R2-DEST destination vector. The hyperactive PB plasmid, pCMV-hyperactive PBase, was kindly provided by the Medical Research Council^23^. This vector was linearized via digestion with AgeI, and the mRNA was synthesized using mMESSAGE mMACHINE (tm) T7 ULTRA Transcription Kit (Thermo Fisher Scientific Inc.). The mRNA was purified using a MEGAclear Kit (Thermo Fisher Scientific Inc.). For microinjection, a PB vector (0.66 ng/μL or 2 ng/μL) and transposase mRNA (0.33 ng/μL or 1 ng/μL) were diluted with saline to obtain final concentrations of 1 ng/μL and three ng/μL, respectively.

### Production of Tg offspring in marmoset

Ovarian cycle control was performed as previously reported^4^. The ovarian cycles of luteal phase adult females were reset via intramuscular administration of 0.8 µg prostaglandin F2α (PGF2α, Estrumate; MSD Animal Health, Tokyo, Japan) to induce luteolysis, and the onset of the follicular phase was confirmed based on blood progesterone levels the day after PGF2α injection. Serum progesterone levels were measured using an enzyme immunoassay kit (TOSOH, Tokyo, Japan), and ovulation was defined the day before the serum progesterone concentration exceeded 10 ng/mL.

Marmoset follicle development was stimulated via intramuscular injection of human follicle-stimulating hormone (hFSH; 25 international units (IU); Fuji Pharmaceuticals, Tokyo, Japan). hFSH administration was performed every other day for 9 days, and human chorionic gonadotropin (hCG; 75 IU; ASKA Pharmaceuticals, Inc. Ltd., Tokyo, Japan) was administered at 17:00 on the day following the ninth FSH administration. After 8 h of hCG administration, oocytes were collected via laparotomy. Anesthesia was performed using the same protocol as previously reported^4^. Briefly, female marmosets received 0.04 mg/kg of medetomidine (Domitor; Nippon Zenyaku Kogyo, Fukushima, Japan), 0.40 mg/kg of midazolam (Dormicam; Astellas Pharma, Tokyo, Japan), and 0.40 mg/kg of butorphanol (Vetorphale Meiji Seika Pharma, Tokyo, Japan) as preanesthetic. During surgery, marmosets were anesthetized through isoflurane (DS Pharma, Osaka, Japan) inhalation. Postoperative medication including 1.2 mg/kg ketoprofen (Capisten; Kissei Pharmaceutical, Matsumoto, Japan) as analgesic and 15 mg/kg ampicillin (Viccillin; Meiji Seika Pharma Co., Ltd., Tokyo, Japan) as antibiotic was administered once daily for more than two consecutive days.

Oocytes were collected in porcine oocyte medium (Research Institute for the Functional Peptides, Yamagata, Japan) supplemented with 5% heat-inactivated FBS (Thermo Fisher Scientific Inc.) and 75 IU/mL FSH (Fuji Pharmaceuticals, Tokyo, Japan), and *in* vitro maturation (IVM) was performed for 26 h. For *in vitro* fertilization, marmoset semen was collected via penile vibratory stimulation^29,40^. The semen was washed with TYH medium (LSI Medience Corporation, Tokyo, Japan), and the sperm suspension was incubated in TYH medium for 30 min to allow the sperm to swim up. After incubation, the sperm suspension was adjusted to a final sperm concentration of 3.6 × 10^6^ sperms/mL. Matured oocytes were inseminated with sperm suspension for 18 h, and fertilized oocytes were retrieved by cytoplasmic injection. Circular PB vector and hyPBase mRNA were injected into the cytoplasm. Injected oocytes were subjected to in vitro culture in sequential Cleave (CooperSurgical, Inc., Trumbull, CT, USA) until the 8-cell stage. They were cultured in sequential Blast (CooperSurgical, Inc.) for the remaining period up to the blastocyst stage. IVMFC was performed via incubation under a gas phase of 5% CO^2^, 5% O^2^, and 90% N^2^ at 38.0 ºC.

Fluorescence protein expression was confirmed in the developed embryos using inverted fluorescence microscopy, and the embryos expressing EGFP only were selected for embryo transfers. For embryo transfers, the ovarian cycles of donor and recipient animals were synchronized via PGF2α administration, and ovulation of the recipient animals was determined by plasma progesterone levels exceeding 10 ng/mL. Non-surgical embryo transfer was performed after 3–5 days of ovulation of recipient animals. The transfer method was performed as previously reported^29^.

### Germline transmission of the transgene

Sperms were collected from sexually mature male Tg marmosets to study germ line transmission as previously described^4^. Fertilized oocytes were obtained via *in vitro* fertilization as described above. Alternatively, embryos were collected by spontaneous mating with WT and non-surgical uterine flushing. Embryo collection from the uterus was performed as previously described^41^. Anesthesia was applied as described above. A blunt-end stainless steel stylet (25 G, 120 mm-long) covered with an 18 G, 108 mm-long Fluon ETFE catheter (Oviraptor, Altair, Yokohama, Kanagawa, Japan) was inserted into the uterus through the cervix, and only the blunt-end stainless steel stylet was removed. The flushing medium, Waymouth’s Medium (Thermo Fisher Scientific Inc.), 10% inactivated FBS (Biowest, Nuaille, France), and 1 M HEPES (Sigma-Aldrich, St Louis, MO, USA) were filled in a polyethylene cannula (length 160 mm, inner diameter 0.28 mm, outer diameter 0.61 mm) (Althea, Yokohama, Kanagawa, Japan), which was inserted into the uterus through a Fluon ETFE catheter (Oviraptor, Althea) for flushing. During flushing, the oviduct was compressed through the abdominal wall to prevent the embryos from being released into the abdominal cavity. The flushing medium was collected into Petri dishes through a Fluon ETFE catheter to obtain embryos. The resulting embryos were observed for fluorescence expression under a fluorescence microscope, and EGFP-positive embryos were transferred as above.

### Statistical analysis

To evaluate differences between experimental groups, a χ^2^-test was performed. Differences at p *<* 0.05 were considered statistically significant.

## Acknowledgements

We thank all members of the Department of Marmoset Biology and Medicine and Center of Basic Technology in Marmoset for maintenance of animals at the Central Institute for Experimental Animals. We thank Dr. Kanki for providing pENTR-hKOd2PEST plasmid.

## Funding

The study was supported by grants from the Strategic Research Program for Brain Sciences from Japan Agency for Medical Research and development (AMED) under Grant Numbers JP17dm0107051 and JP18dm0207065 and was partially supported by Grants-in-Aid for Scientific Research (MEXT/JSPS KAKENHI Grant Number 17K14979).

## Author contributions

E.S., H.O., T.T., and T.M. designed the study. T.M., W.K., and T.T. designed and constructed vectors. M.G., T.T., and E.T. assisted in mouse embryological technique development. T.T. and M. Y. contributed to genotyping. R.I. conducted FACS analysis. E.S. and T.T. wrote the manuscript. All authors revised and edited the manuscript.

## Competing interests

The authors declare no competing interests.

## Data availability statement

All data generated or analyzed during this study are included in this published article (and its Supplementary Information files).

## Notes

### Competing Interest Statement

The authors have declared no competing interest.

