## Supplementary methods for "Strategy for selective acquisition of transgenic marmosets using the piggyBac transposon system"

### Cell Transfection

HEK293T cells were cultured in complete Dulbecco's Modified Eagle Medium supplemented with 10% heat-inactivated fetal bovine serum (FBS, Thermo Fisher Scientific Inc.). Cells were seeded on 24-well plates at $1\times10^5$ cells per well. Following this, 400 ng of PB-plasmid and 200 ng of hyPBase-plasmid were transfected using Lipofectamine LTX (Thermo Fisher Scientific Inc.). The cells were observed via fluorescence microscopy, and they were analyzed for EGFP or hKO1-positive cells via FACS.

### Production of Tg mice

Mouse zygotes for genetic modification were prepared via in vitro fertilization as previously described[1]and they were vitrified approximately 8 h after insemination[1] Microinjection was performed after 1 hour of warming the vitrified zygotes[2]. The microinjected zygotes were cultured with KSOM medium under 5% CO2, 5% O2, and 90% N2 at 37 °C, and they were observed via fluorescence microscopy every day until the embryos reached the blastocyst stage. The solutions used for embryo manipulations were purchased from Arc Resources (Kumamoto, Japan). The obtained blastocyst was transplanted into the uterus of a pseudo-pregnant female mouse[3], and the obtained Tg mice were mated with a wild-type mouse to produce the next generation.

### Chromosome preparation and Fluorescent in situ hybridization

Spleen cells were cultured with RPMI1640 medium supplemented with 20% FBS, 3 µg/mL concanavalin A, and 10 µg/mL lipopolysaccharide for 2 days. Following this, 300 µg/mL thymidine was added to the medium, and cells were cultured for another 16 h. After changing to flesh medium, 30 µg/mL 5-bromo-2'-deoxyuridine was added for 3.5 h, and following this, 0.02 µg/mL colcemid was added for another 30 min before harvesting. Cultured cells were then treated with 0.075 M KCl for 20 min, and they were washed 3 times in fixative solution (methanol:acetic acid=3:1) before spreading the cells on slides. After drying the slides overnight, the metaphase spreads were stained with Hoechst 33258, and UV irradiation was performed at 70 ºC for 4 min to make chromosome banding patterns.

EGFP and hKO1 were labeled via nick translation with Cy3-dUTP. The labeled probes were mixed with sonicated salmon sperm DNA and mouse cot-1 DNA in hybridization solution. The probe was applied to metaphase spreads, denatured at 70 ºC for 5 min, and hybridized at 37 ºC overnight. The hybridized slide was washed and mounted with antifade mountant. FISH images were captured with the CW4000 FISH application program (Leica

Microsystems Imaging Solution Ltd.) using a cooled CCD camera mounted on a Leica DMRA2 microscope.

#### Genotyping

Tg offspring were identified via genomic PCR or RT-PCR. Genomic DNA was extracted from tissue samples using the DNeasy kit (Qiagen, Hilden, Germany). To determine transgene expression, total RNA was extracted (Takara bio, Shiga, Japan) and reverse-transcribed using ThermoScript RT-PCR Systems (Thermo Fisher scientific Inc.). To detect gene expression, EGFP and hKO1 primers were used. As internal controls, actin was used in marmosets, and the intron of chromosome 15 was used in mice. Primer sequences are shown in Supplementary Table S4. Genomic PCR was performed with the following conditions: 94 °C for 2 min; 31 cycles at 94 °C for 10 s, 63 °C for 30 s, and 68 °C for 20 s. RT-PCR was performed at the following conditions: 94°C for 2 min; 30 cycles at 94°C for 10 s, 63°C for 30 s, and 68°C for 20 s.

### Immunohistochemical analysis

Marmoset placentas were fixed in 10% neutralized formalin. Formalin-fixed tissues were embedded in paraffin, and they were analyzed via hematoxylin-eosin staining and immunohistochemistry. Staining of sections with rabbit polyclonal anti-GFP (Abcam plc, Cambridge, UK) was performed on a fully automated BOND-MAX system (Leica Biosystems, Mount Waverley, VIC, Australia).
