## Supplementary_Figures_and_Tables for "Strategy for selective acquisition of transgenic marmosets using the piggyBac transposon system"

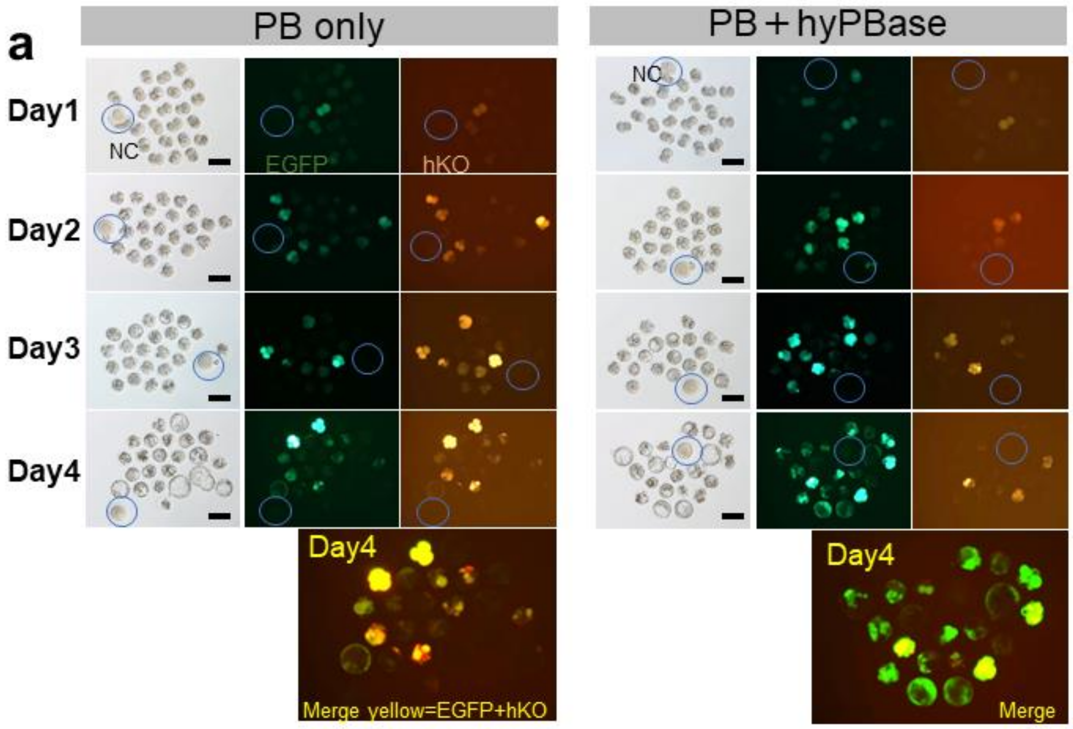


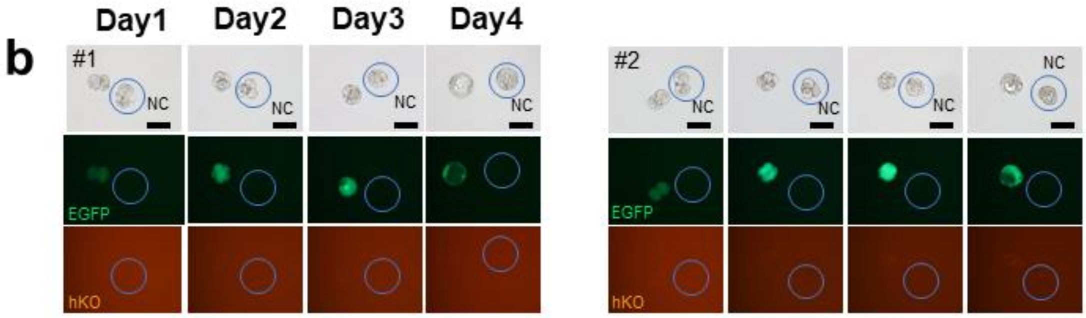


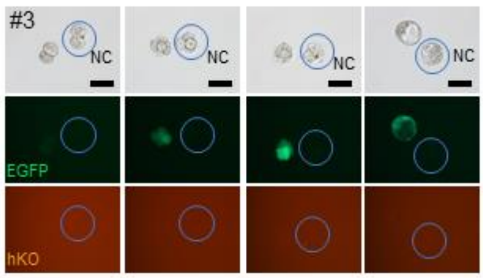


Fig S1. Changes in fluorescence protein expression over time after microinjection in mouse embryos. (a) Left panel: PB only (no transposase), right panel: PB+hyPBase; fluorescence change with transposase-enhanced integration. Bottom panel: merged images of hKO1 and EGFP. The overlapping expression of both is shown in yellow. (b) Changes in fluorescent protein expression over time in a single embryo. Scale bar = 100 μm.

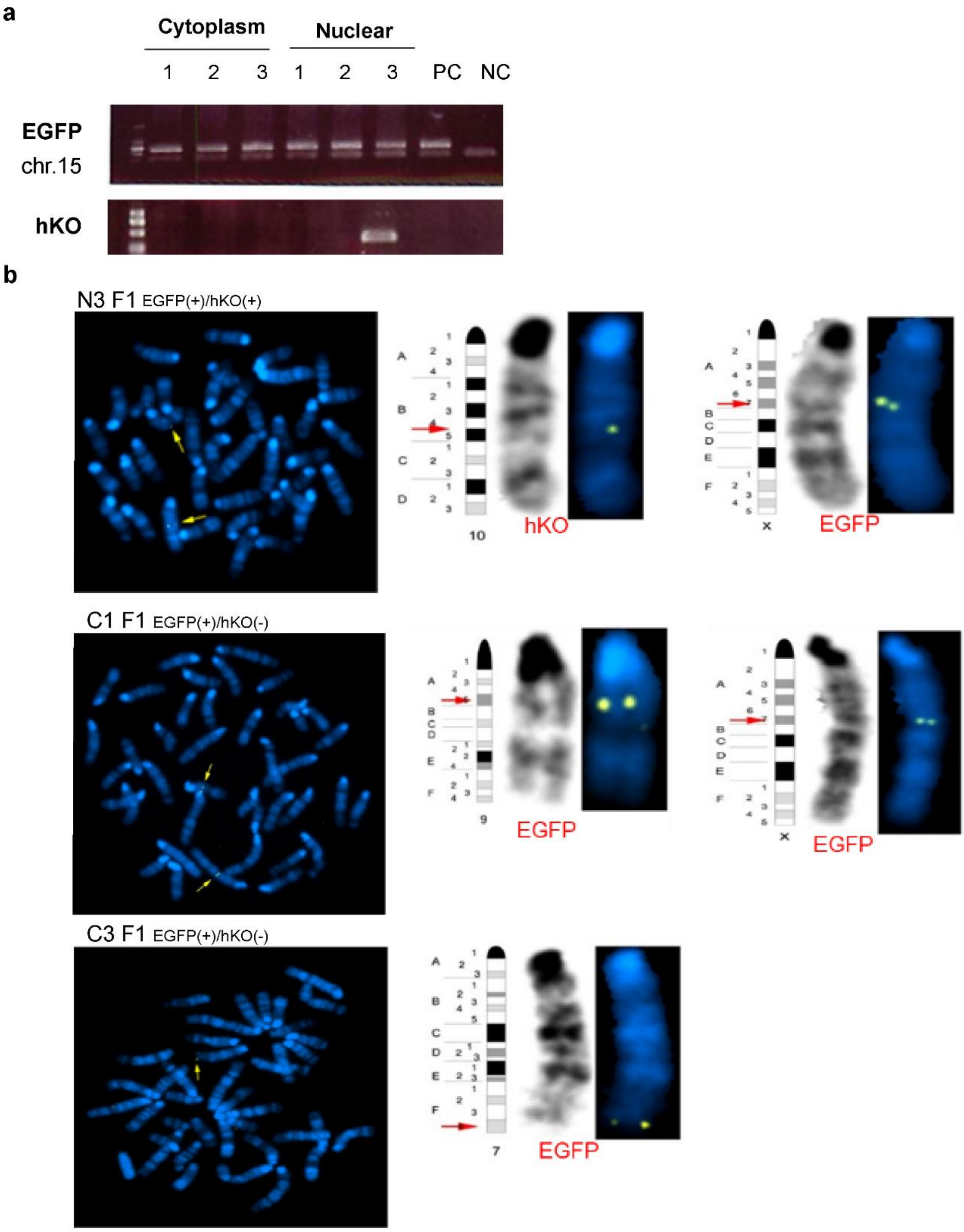


Fig S2. (a) Results of Genomic PCR of F0 mouse. PCR products of EGFP (upper bands) and Chr.15 intron (lower band) as the internal control are indicated. (b) FISH analysis of F1 mice. N3F1: F1 animal from mouse no. 3 of the nuclear injection. The hKO1 gene was inserted into chromosome 10, and the EGFP gene was located on the X chromosome. C1F1:F1 animal from mouse no. 1 that was created via cytoplasmic injection. The EGFP gene was inserted into two locations, chromosomes 9 and X. C3F1: F1 animal from no.3 created via cytoplasmic injection. The EGFP gene was inserted into chromosome 7.

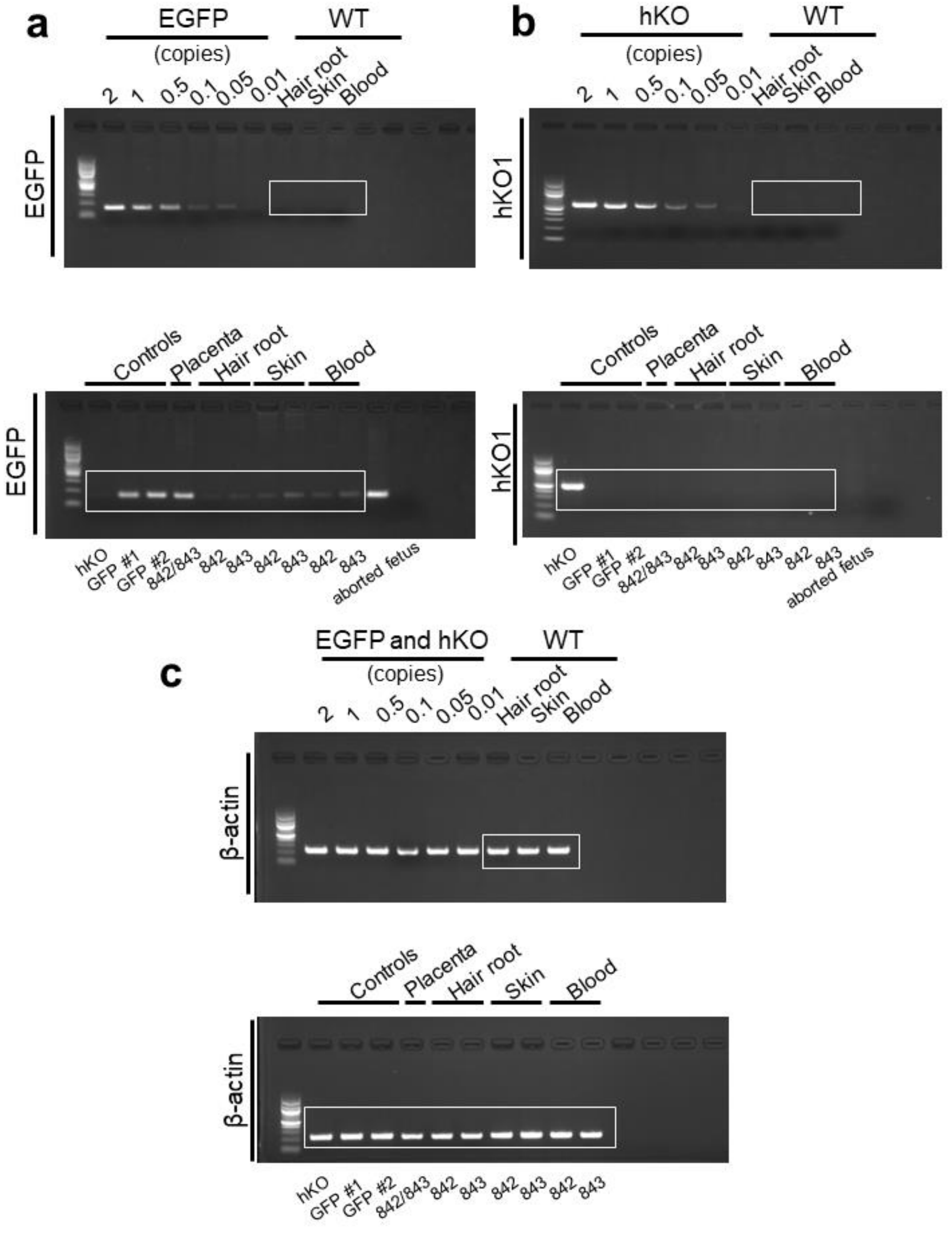


Fig S3. Results of genotyping analysis of Tg marmoset by genomic PCR. This figure complements the cropped gel in the PCR analysis shown in Figure 3d. (a) Results of PCR using GFP primers. (b) Results of PCR using hKO primers. (c) Results of PCR using β-actin. As the control, genomic DNA extracted from WT marmoset fibroblast cells and PB vector were mixed to make controls for 2, 1, 0.5, 0.1, and 0.05 copies of EGFP or hKO, respectively, and were used as the PCR templates. Lanes under the WT label and lanes 842 and 843 indicate results of PCR using genomic DNA extracted from hair roots, skin, and blood as the templates. White boxes indicate a part of the gel shown in the main Figure 3d.

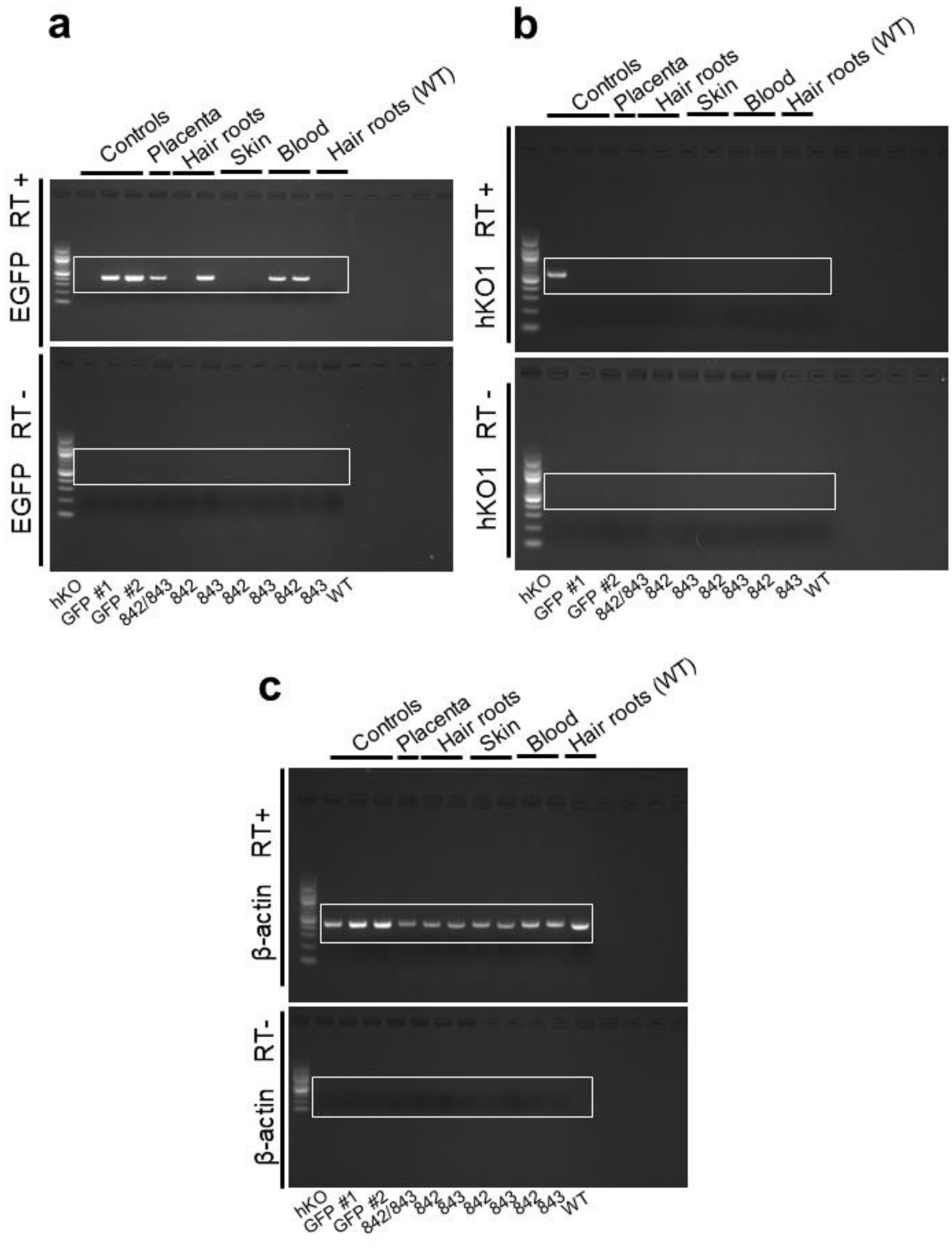


Fig S4. Results of genotyping analysis of Tg marmoset by RT-PCR. This figure complements the cropped gel in the PCR analysis shown in Figure 3e. (a) Results of RT-PCR using EGFP primers. (b) Results of RT-PCR using hKO primers. (b) Results of RT-PCR using β-actin primers.

As the controls, mRNAs extracted from hair roots of marmosets carrying the EGFP or hKO genes were used. Results of RT-PCR from each animal's tissue, hair roots, skin, and blood from 842 and 843 are indicated next to the control lanes. Each upper gel indicates RT-PCR products, and the lower gel show PCR amplicons without reverse transcriptase as the control. White boxes indicate a part of the gel shown in the main Figure 3e.

| Injection | concentration total (PB: hyPBase ) | Color selection | No.of embryo used | Blastocyst (% embryo) | hKO (-) and EGFP (+) Blastocyst (% Blastcyst) | hKO (+) and EGFP (+) Blastocyst (% Blastcyst) | No. of embryos transferred | No.of fetuses (% transfered) | No.of Weanings | No. of EGFP (+) Tg offspring (% Weanings) | No. of hKO (+) Tg offspring (% Weanings) |
|---|---|---|---|---|---|---|---|---|---|---|---|
| Nuclear | 1ng/μl (0.67ng/μl:0.33ng/μl) | - | 70 | 51 (72.9) | - | - | 51 | 16 (31.4)$^{a}$ | 8 | 4 (50) | 0 (0) |
| | | + | 92 | 66 (71.7) | 55 (83.3) $^{a}$ | 0 (0) | 39 | 5 (12.8)$^{b}$ | 3 | 3 (100) | 1 (33.3) |
| Cytoplasm | 3ng/μl (2ng/μl:1ng/μl) | - | 74 | 48 (64.9) | - | - | 48 | 14 (29.2)$^{ab}$ | 11 | 9 (81.8) | 1 (9.1) |
| | | + | 95 | 64 (67.4) | 27 (42.2)$^{b}$ | 0 (0) | 26 | 3 (11.5)$^{ab}$ | 3 | 3 (100) | 0 (0) |

Table S1. Comparison of the birth rates of Tg animals via fluorescent protein selection after PB microinjection in mouse.

Values within the same column with different letters (a, b) differ significantly ($p<0.05$), $\chi 2$-test.

| Transgenenesis method (PB plasmid: hyPBase mRNA) | Name of Founders | No. of F1 offspring | No. of EGFP (+) F1 offspring (% per offspring) | No. of hKO1 (+) F1 offspring (% per offspring) |
|---|---|---|---|---|
| Nuclear 1ng/μl (0.67ng/μ:0.33ng/μ) | N1 | 0 | 0 | 0 |
| | N2 | 10 | 0 | 0 |
| | N3 | 6 | 6 (100) | 4 (66.7) |
| Cytoplasm 3ng/μl (2ng/μ:1ng/μ) | C1 | 9 | 2 (22.2) | 0 |
| | C2 | 8 | 5 (62.5) | 0 |
| | C3 | 8 | 3 (37.5) | 0 |

Table S2. Germline transmission of Tg in first fillia (F1) offspring from founder x WT.

| Animal | Primer name | Primer Sequence (Forward) | Primer Sequence (Reverse) |
|---|---|---|---|
| Mouse | EGFP | CTGGTCGAGCTGGACGGCGAC | CACGAACTCCAGCAGGACCATG |
| | hKO1 | AAGCCCGAGATGAAGATGAA | ATCTCCTCGGGGTACTTGGT |
| | intron Chr 15 (internal control) | AATAGCAAACTCACATGATCCTTGGC | ACTGCCTTGAACAACATTCCCTG |
| Marmoset | EGFP (genomic) | CACAACGTCTATATCATGGC | TGCTCAGGTAGTGGTTGT |
| | hKO1 (genomic) | CAAGCCCGAGATGAAGATG | GGCATCTTCAGGATCTTCTT |
| | β－actin (internal control:genomic) | TGTAGGTACTAACACTGGCTCGTGTGACAA | GGGTGTTGAAGGTCTCAAACATGATCTGTA |
| | EGFP (RT) | CAAGGACGACGGCAACTACAAGACC | GCTCGTCCATGCCGAGAGTGA |
| | hKO1 (RT) | GCTCGTCCATGCCGAGAGTGA | GGCATCTTCAGGATCTTCTT |
| | β－actin (internal control: RT) | TGGACTTCGAGCAGGAGAT | CCTGCTTGCTGATCCACATG |

Table S3. List of PCR primer sequences.
